# Pyruvate carboxylase supports the pulmonary tropism of metastatic breast cancer

**DOI:** 10.1101/317743

**Authors:** Aparna Shinde, Tomasz Wilmanski, Hao Chen, Dorothy Teegarden, Michael K. Wendt

## Abstract

**Background:** Overcoming systemic dormancy and initiating secondary tumor grow under unique microenvironmental conditions is a major rate-limiting step in metastatic progression. Disseminated tumor cells encounter major changes in nutrient supplies and oxidative stresses compared to the primary tumor and must demonstrate significant metabolic plasticity to adapt to specific metastatic sites. Recent studies suggest that differential utilization of pyruvate sits as a critical node in determining the organotropism of metastatic breast cancer. Pyruvate carboxylase (PC) is key enzyme that converts pyruvate into oxaloacetate for utilization in gluconeogenesis and replenishment of the TCA cycle.

**Methods:** Patient survival was analyzed with respect to gene copy number alterations and differential mRNA expression levels of PC. Expression of PC was analyzed in the MCF-10A, D2-HAN and the 4T1 breast cancer progression series under *in vitro* and *in vivo* growth conditions. PC expression was depleted via shRNAs and the impact on *in vitro* cell growth, mammary fat pad tumor growth, and pulmonary and non-pulmonary metastasis was assessed by bioluminescent imaging. Changes in glycolytic capacity, oxygen consumption and response to oxidative stress were quantified upon PC depletion.

**Results:** Genomic copy number increases in *PC* were observed in 16-30% of metastatic breast cancer patients. High expression of PC mRNA was associated with decreased patient survival in the MCTI and METABRIC patient datasets. Enhanced expression of PC was not recapitulated in breast cancer progression models when analyzed under glucose-rich *in vitro* culture conditions. In contrast, PC expression was dramatically enhanced upon glucose deprivation and *in vivo* in pulmonary metastases. Depletion of PC led to a dramatic decrease in 4T1 pulmonary metastasis but did not affect orthotopic primary tumor growth. Tail vein inoculations confirmed the role of PC in facilitating pulmonary, but not extra-pulmonary tumor initiation. PC-depleted cells demonstrated a decrease in glycolytic capacity and oxygen consumption rates and an enhanced sensitivity to oxidative stress.

**Conclusions:** Our studies indicate that PC is specifically required for the growth of breast cancer that has disseminated to the lungs. Overall, these findings point to the potential of targeting PC for the treatment of pulmonary metastatic breast cancer.

## Background

Metastasis of primary mammary tumors to vital secondary organs is the primary cause of breast cancer-associated death, with no effective treatment [1]. Metastasis is a highly selective process that requires cancer cells to overcome multiple barriers to escape the primary tumor, survive in circulation, and eventually colonize distant secondary organs. The major sites of breast cancer metastasis include bone, liver, brain and lungs [2]. Understanding the unique drivers of organ-specific metastases could hold the key to more personalized and more effective therapies for stage 4 patients. Previous studies have sought to characterize stable genetic and gene expression changes that drive metastatic tropism of individual cancer cells [3,4]. However, the plasticity of metastatic breast cancer cells in response to specific growth environments makes identification of *in vivo* tropic factors difficult.

Metabolic reprogramming is a hallmark of cancer, and is implicated in cell proliferation, survival and metastatic progression [5]. Considerable progress has been made in understanding the unique metabolic changes cancer cells must undergo to adapt to the hypoxic and acidic microenvironment of the primary tumor [6]. Furthermore, the Warburg effect dictates that tumor cells will continue to utilize glycolysis for energy production even in the presence of abundant oxygen. However, recent studies suggest that metastatic cells have an increased ability to alternate between usage of glycolysis or oxidative phosphorylation (OXPHOS) for energy production in response to particular environmental stresses [7]. Furthermore, expression of the master transcriptional regulator of mitochondrial biogenesis, peroxisome proliferator-activated receptor gamma coactivator 1-alpha (PGC-1α), has recently been linked to the pulmonary metastasis of breast cancer [8]. Overall, these and other studies support the notion that breast cancers capable of reversing the Warburg effect and reactivating mitochondrial OXPHOS are at a selective advantage during initiation of metastatic tumor growth, particularly within the oxygen-rich pulmonary microenvironment.

The axis of pyruvate utilization is emerging as a key regulatory point in cancer cell metabolism, and may be critical to organotropism of metastatic breast cancer [9–11]. The majority of pyruvate in primary tumors is converted into lactate to sustain high rates of glycolysis. Indeed, depletion of lactate dehydrogenase (LDH), the enzyme responsible for lactate production inhibited primary tumor growth and subsequently metastasis [12]. Alternatively, pyruvate may enter the TCA cycle via its metabolism by pyruvate dehydrogenase (PDH) or pyruvate carboxylase (PC). Dupuy *et al.* demonstrated that a key negative regulator of PDH, pyruvate dehydrogenase kinase (PDK1), is essential for liver metastasis [13]. These data suggest that modulation of mitochondrial pyruvate metabolism may determine successful colonization of specific organs by metastatic cancer cells. PC is a mitochondrial enzyme which sustains anaplerosis through its carboxylation of pyruvate into oxaloacetate. PC has recently been shown to be upregulated in lung metastases [8]. Given these data and the critical role of pyruvate metabolism in metastasis, we sought to address the hypothesis that PC is specifically required for the initiation of metastatic outgrowth within the pulmonary microenvironment.

The current study characterizes PC expression and genomic amplification in relation to breast cancer patient survival. Furthermore, we utilize several models of breast cancer to delineate the pulmonary tropism that is dictated by PC. Overall, our data clearly indicate that PC is specifically required for initiation of pulmonary metastatic breast cancer, but it is not required for extra-pulmonary tumor growth. These findings suggest inhibition of PC may serve as an effective therapeutic target for the treatment of pulmonary metastasis.

## Methods

### Cell lines and reagents

The five different TRC lentiviral mouse *Pcx*-targeting shRNAs (lentiviral pLKO.1 TRC cloning vector) were purchased from GE Dharmacon. The shRNA lentiviral plasmids were cotransfected with psPAX2 and pMD2.G into HEK293T cells using polyethylenimine to obtain lentiviral particles. The 4T1, 4T07 and D2.A1 cells were transduced with lentiviral particles for 48 hours and stably transduced cells were selected over a span of 14 days in puromycin (5μg/ml). In all cases, separate cells were transduced with scrambled non-silencing shRNAs as a control. The target shRNA sequences were 5’- AAAGGACAAATAGCTGAAGGG-3’(shPC 25) and 5’ – TTGACCTCGATGAAGTAGTGC-3’ (shPC28). The scrambled sequence was 5’-TTCTCCGAACGTGTCACGT-3’. All cells were culture in DMEM containing, 10% fetal bovine serum, Pen/Strep, 25 mM glucose, 1 mM Sodium Pyruvate and 4 mM Glutamine. Where indicated glucose concentrations were decreased to 5.6 mM.

### Immunological assays

For immunoblot analyses cells were lysed using a modified RIPA lysis buffer containing 50 mM Tris, 150 mM NaCl, 0.25% sodium deoxycholate, 1.0% NP40, 0.1% SDS, protease inhibitor cocktail, 10 mM activated sodium ortho-vanadate, 40 mM β-glycerolphosphate and 20 mM sodium fluoride. These lysates were separated by reducing SDS PAGE and probed for PC (Santa Cruz Biotechnology, Santa Cruz, CA), actin (Santa Cruz Biotechnology, Santa Cruz, CA), or β-tubulin (DSHB, Iowa City, IA). Immunohistochemical analyses of formalin fixed paraffin embedded tissue sections from 4T1 primary and metastatic tumors were conducted by deparfinization of in xylene, rehydration, and antigen retrieval using 10mM sodium citrate (pH 6.0) under pressurized boiling. After inactivation of endogenous peroxidases in 3% H_2_O_2_, primary antibodies specific for PC (Sigma, St. Louis, MO) or HIF1-α (Novus, Littleton, CO) were added and incubated overnight. Protein specific staining was detected through the use of appropriate biotinylated secondary antibodies in conjunction with ABC reagent (Vector, Burlingame, CA). These sections were counterstained with hematoxylin, dehydrated, and mounted.

### 3D culture

Bioluminescent 4T1 scram, shPC 25 and shPC 28 cells were grown under 3D culture conditions. Cell growth was quantified via addition of luciferin (GoldBio, St. Louis, MO). Briefly, 1000 cells were plated in each well of a white-walled 96-well dish on top of a solidified 50 μl bed of Cultrex basement membrane extract (BME) from (Trevigen, Gaithersburg, MD). These cells were suspended in growth media containing DMEM (low or high glucose), 10% FBS and 5% of the BME.

### Invasion and migration assays

Control and PC depleted 4T1 cells were plated at equal cell density in serum free medium containing either 1 or 4 mM pyruvate into 8μm fluoroBlok 24-well transwell inserts (Corning, NY). The bottom well was filled with 10% FBS medium containing either 1 mM or 4 mM pyruvate. Serum free medium in the bottom well was used as a negative control. Cell migration was quantified after 12 hours of incubation using 5 μg/mL of Calcein AM in PBS and a Synergy H1 Multi-mode reader (bottom read: ex./em. 495/530). For wound closure assays control and PC depleted 4T1 cells were plated onto 6-well plates and grown till 90-95% confluence. Once nearly confluent, each well was scratched using a 200ul pipette tip and cell media was replaced to 0.5% serum. Wound images were taken at time 0,15 and 24 hours. Open wound area was quantified using TScratch software [15] and percent wound closure was calculated using the following formula: [(open area at time 0 - open area at time x)/open area at time 0] *100.

### Metabolic and oxidative stress analysis

Rate of glycolysis and oxygen consumption were analyzed using the Seahorse XFe24 Analyzer (Agilent, Santa Clara, CA). 4T1 shScram and shPC25 cells were plated at a concentration of 20,000 cells/well on a Seahorse bioanalyzer plate 24 hours before analysis. 4T07 shScram and shPC25 cells were plated at a concentration of 40,000 cells/well on the same Seahorse bioanalyzer plate to achieve comparable viable cell number at the time of experiment. The glycolytic stress test was conducted according to the manufacturer’s instructions with 10 mM glucose concentration in the media while oxygen consumption rate (OCR) was measured in the presence of 10mM glucose, 2 mM glutamine, and 1 mM pyruvate. After the assay was completed, cells within each well were lysed into 25 ul of 1x RIPA buffer and protein content was measured using a BCA Assay. Where indicated 4T1 cells were plated overnight and treated with hydrogen peroxide (H_2_O_2_) at multiple concentrations on day 2. The cells were washed with PBS twice on day 7, and stained with crystal violet, lysed and absorbance was read at 600nm.

### *In Vivo* Studies

Control or PC-depleted 4T1 cells previously engineered to express firefly luciferase stably were resuspended in sterile PBS (50 μl) and injected orthotopically into the mammary fat-pad (2.5 × 10^4^ cells/mouse) of 6-week-old female Balb/c mice. Primary tumor growth and metastasis development was assessed by using digital calipers and by weekly bioluminescent imaging on an advanced molecular imager (AMI, Spectral Instruments; Tucson AZ). For tail vein assays, control or PC depleted 4T07 or D2.A1 cells previously engineered to express firefly luciferase were resuspended in sterile PBS (200 μl) and injected into lateral tail vein (5 × 10^5^ cells/injection) of 6-week-old female Balb/c mice. Mice were assessed for tumor development by weekly bioluminescent imaging on a AMI *in vivo* imager. All animal studies were performed in accordance with the animal protocol procedures approved by the Purdue Animal Care and Use Committee of Purdue University.

### Statistical analyses

One way ANOVA or 2-sided T-tests were used where the data met the assumptions of these tests and the variance was similar between the two groups being compared. Expression values for *PC* across breast cancer subtypes were analyzed using a Kruskal-Wallis test, together with a Dunn’s multiple comparisons test. P values of less than 0.05 were considered significant. No exclusion criteria were utilized in these studies.

## Results

### PC expression is increased in more aggressive breast cancers

To evaluate the importance of PC in breast cancer progression we initially utilized the BreastMark Kaplan-Meier analysis program to analyze the Molecular Therapeutics for Cancer, Ireland (MTCI) dataset [16]. There was a significant decrease in survival of patients whose tumors express high versus low-levels of PC based on the mean value of the entire cohort (Fig. 1a). We also analyzed the Molecular Taxonomy of Breast Cancer International Consortium (METABRIC) dataset consisting of 2509 patient samples annotated for several pieces of the patient-specific data including breast cancer subtype and tumor stage [17]. Analysis of PC gene copy number in the METABRIC and other invasive and metastatic breast cancer datasets indicated that 16% to 30% of patients demonstrate copy number gains in PC (Table 1). All patient datasets analyzed and the cell line encyclopedia indicated increased PC gene copy number directly correlates with increased PC mRNA expression levels (not shown). Importantly, patients with PC gene amplification in the METABRIC data set displayed significantly reduced survival times as compared to the rest of the cohort (Fig. 1b). Significant differences in PC mRNA expression were observed between subtypes, with the aggressive basal subtype showing the highest mean level of PC expression (Fig. 1c). Changes in PC mRNA expression was also analyzed across tumor stage. No significant correlatation in PC expression was observed in regard to tumor stage or tumor size (Fig 1d, Figure S1). However, significant outliers were determined by a ROUT analysis in both stage 1 and stage 2 patients (Fig. 1d). The high-level PC outlier patients in the stage 1 group demonstrated a significant decrease in survival as compared to the rest of the stage 1 patients (Fig. 1e). These data suggest that high-level PC expression does not contribute to primary tumor growth, but may be involved in the later stages of breast cancer progression, contributing to decreased patient survival. Further supporting these genomic and mRNA analyses, IHC analysis of lobular and ductal breast carcinoma tissues (n=10) demonstrated the detection of PC expressing tumor cells, something that was not observed in normal mammary epithelium (n=3) (Fig. 1f and Figure S2). Overall, these data clearly demonstrate genomic amplification and increased expression of PC in more aggressive breast cancers. Furthermore, lack of PC correlation with primary tumor stage/size suggests a functional role for this protein in driving the later stages of breast cancer progression.

**Figure 1.**
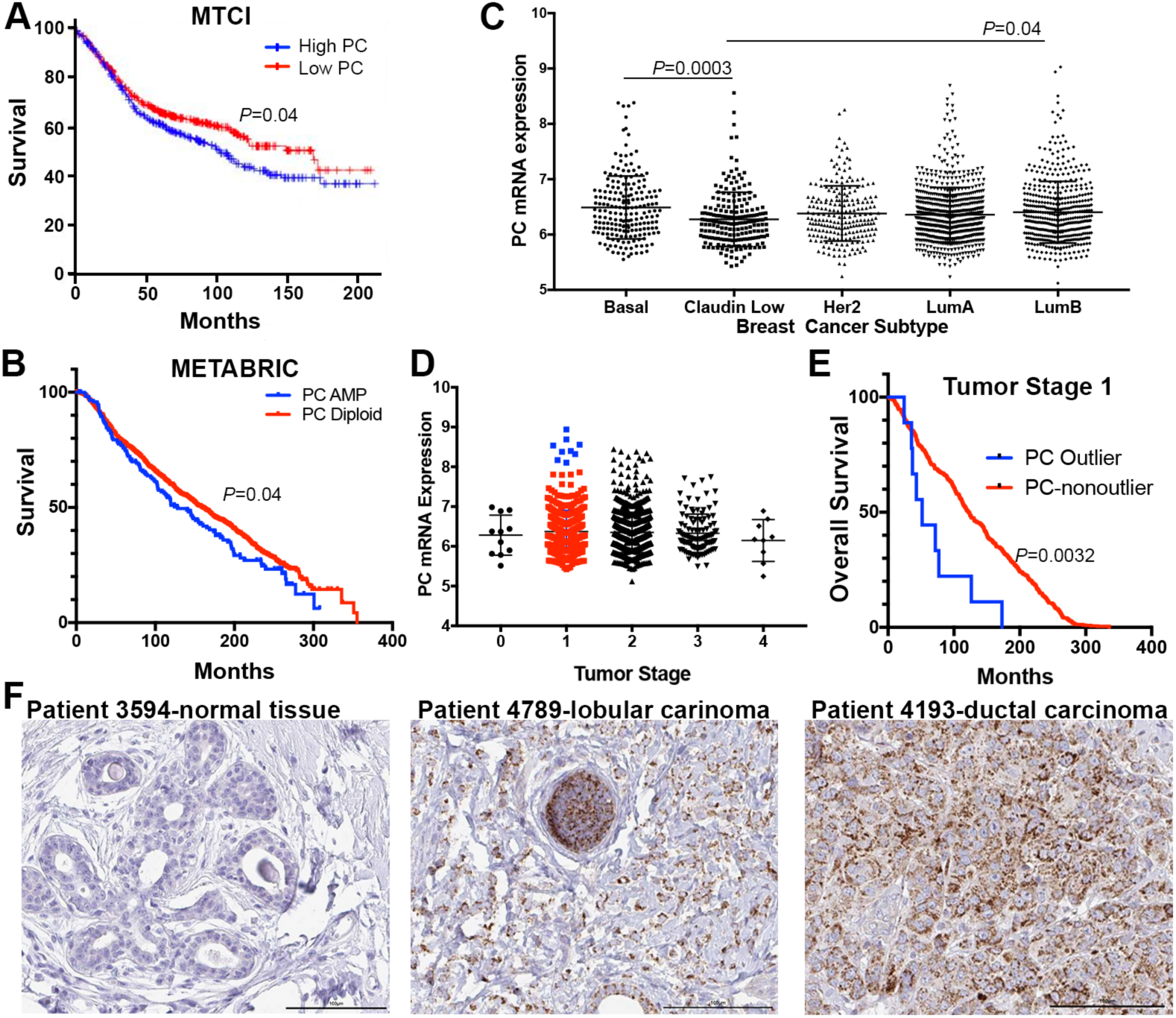
PC expression is increased in more aggressive breast cancers. (A) Patients with lymph node positive breast cancer from the MTCI dataset were split into two groups split based on mean expression of PC and the resultant differences in patient survival are shown. (B) Patients within the METABRIC dataset were separated into two groups based on increased genomic copy numbers of *PC.* Differences in patient survival are shown. (C) Expression levels of PC are plotted across the different breast cancer subtypes within the METABRIC dataset. Subtypes were defined by the PAM50 expression analysis. Differences in PC expression levels were analyzed by a Kruskal-Wallis test resulting in the indicated P values. (D) PC expression levels within the METABRIC data set are plotted based on tumor stage. Significant outliers in the Stage 1 group as determined by a ROUT analysis are shown in blue. (E) The stage 1 patients shown in panel D were separated based on PC outlier status and differences in overall survival are shown. Data in panels A, B and E were analyzed by a Log Rank test resulting in the indicated P values. (F) IHC analyses for PC expression within normal mammary tissue, lobular breast carcinoma and ductal breast carcinoma. Data in panel F are representative of 3 normal samples and 10 primary breast tumors.

**Table 1.**
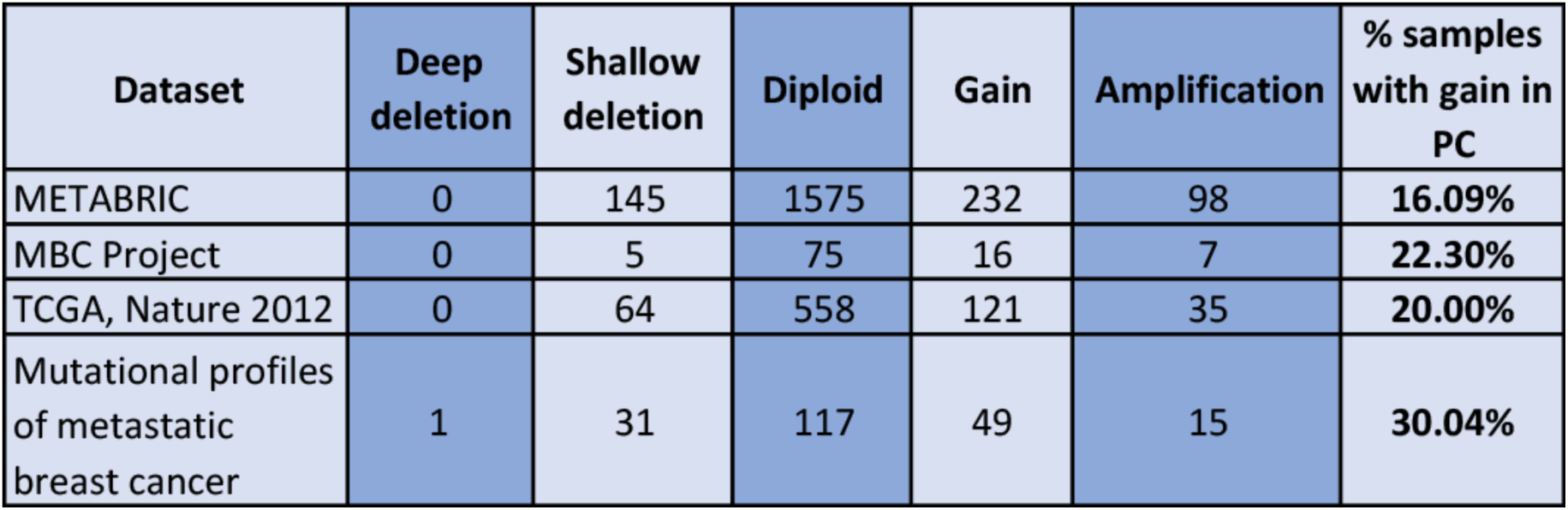
Gene copy numbers of *PC* are increased in metastatic breast cancer patients. Copy number alterations in the *PC* gene (chromosome 11q13.2) were analyzed in the indicated datasets. The percentage of patient tumors demonstrating increases in *PC* gene copy number was calculated for each dataset.

### PC expression is increased in the pulmonary microenvironment

To examine the functional role of PC in breast cancer metastasis we first assessed PC expression levels across several breast cancer progression series. In contrast to the patient data shown in figure 1, PC expression failed to correlate with increasing metastatic capacity of the MCF10A series, the D2-hyperplastic alveolar nodule (HAN) series and the 4T1 series (Fig. 2a) [18–20]. Previous studies suggest that PC expression can be increased during epithelial-mesenchymal transition (EMT) through the transcription factor Snail [9,21]. While stimulation of the 4T1 cells with TGF-β1 induced a robust morphological change consistent with EMT, we did not observe any change in PC expression (Figure S3A and S3B). Furthermore, expression of Snail in the MCF-10-T1k cells did not robustly increase PC expression (Figure S3C). The presence of glucose can enhance expression of PC through activation of one of two distinct promoters [22]. In contrast, PC is also required for gluconeogenesis and therefore its expression can be selectively enhanced under glucose-deprived conditions [23]. We observed that depletion of exogenous glucose from sodium pyruvate containing growth media of 4T1 and 4T07 cells led to increased expression of PC (Fig. 2b). We also observed an increase in PC expression in orthotopic 4T1 primary tumors (Fig. 2c). However, upon IHC analysis we found that PC was highly expressed in the murine sebaceous glands within these tumors, but was still not detectable in 4T1 tumor cells (Fig. 2c and 2d). Expression of PC was further enhanced in 4T1 lung metastases and IHC analyses clearly indicated that PC was expressed in both bronchial epithelial cells and tumor cells (Fig. 2c and 2d; [11]). Taken together these data suggest that in contrast to the primary tumor, growth within the pulmonary microenvironment demands high levels of PC.

**Figure 2.**
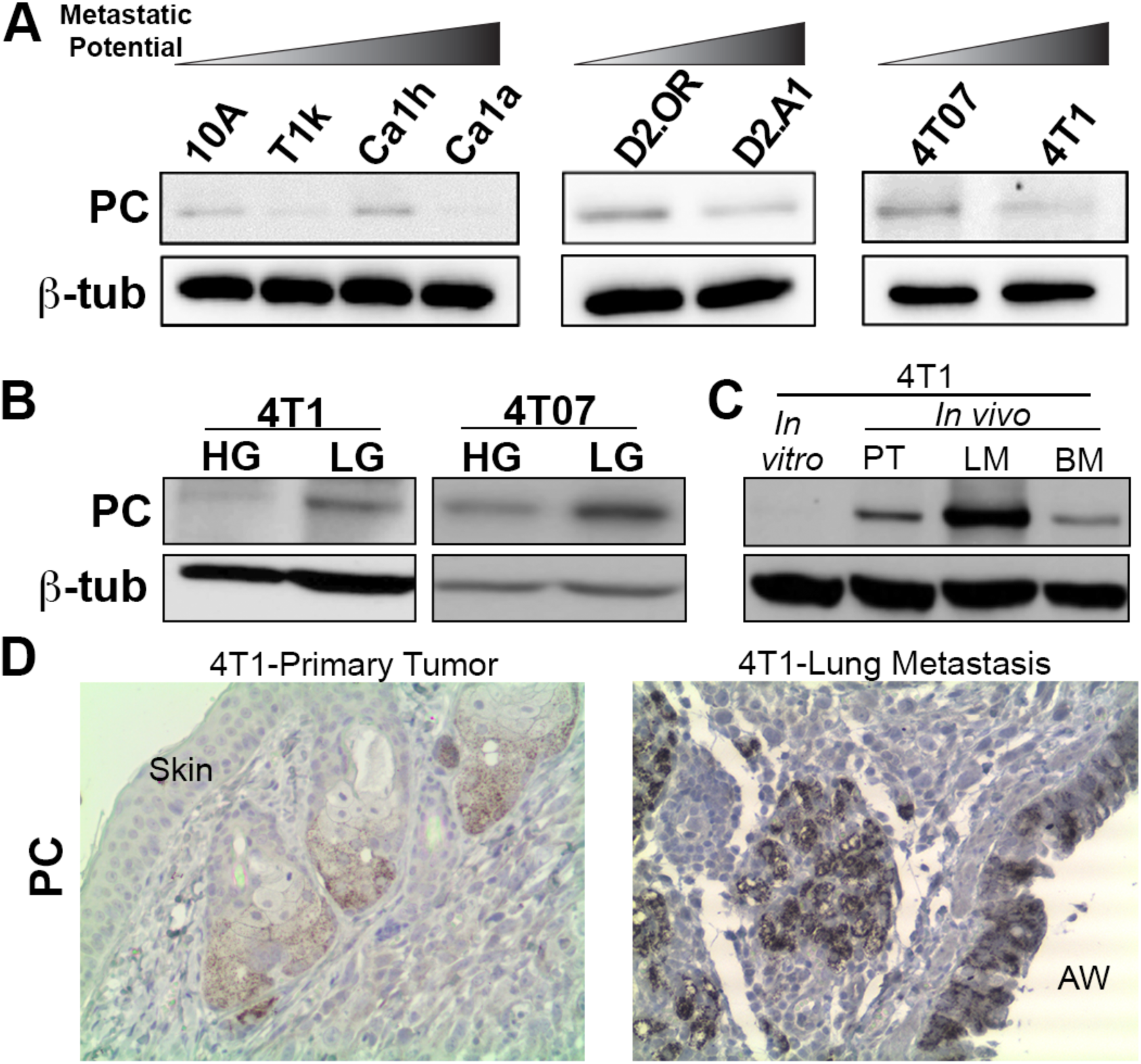
PC expression is enhanced in the pulmonary microenvironment. (A) Expression of PC protein was assessed by immunoblot across the indicated cell lines of varying metastatic potential. (B) 4T07 and 4T1 cells were cultured in the absence of exogenous glucose for 4 days and PC expression was assessed by immunoblot. (C) Expression of PC proteins was analyzed in 4T1 lysates harvested from *in vitro* culture, mammary fat pad primary tumors (PT), and their resultant metastases growing within the lung (LM) and bone marrow (BM). (D) IHC analyses for PC expression within 4T1 primary tumors pushing directly against the skin and the resulting pulmonary metastases growing near an airway (AW) in the lungs. Data are representative of at least 3 tumors from each location. For panel A-C expression of β-tubulin (β-tub) served as a loading control.

### Depletion of PC inhibits spontaneous metastasis

We next depleted PC expression in the highly metastatic 4T1 cells using two independent shRNAs [20]. Depletion of PC did not affect cellular migration as assessed by both a transwell and a wound healing assays (Figure S4). However, depletion of PC did result in inhibition of cell growth irrespective of glucose concentration or whether cells were grown on two-dimensional (2D) plastic or within three-dimensional (3D) hydrogel cultures (Fig. 3). In contrast to *in vitro* culture, there were no significant differences in the growth of PC-depleted primary tumors as compared to control (Fig. 4a and 4b). However, resultant pulmonary metastases observed in these mice were drastically inhibited upon depletion of PC (Fig. 4c-4e).

**Figure 3.**
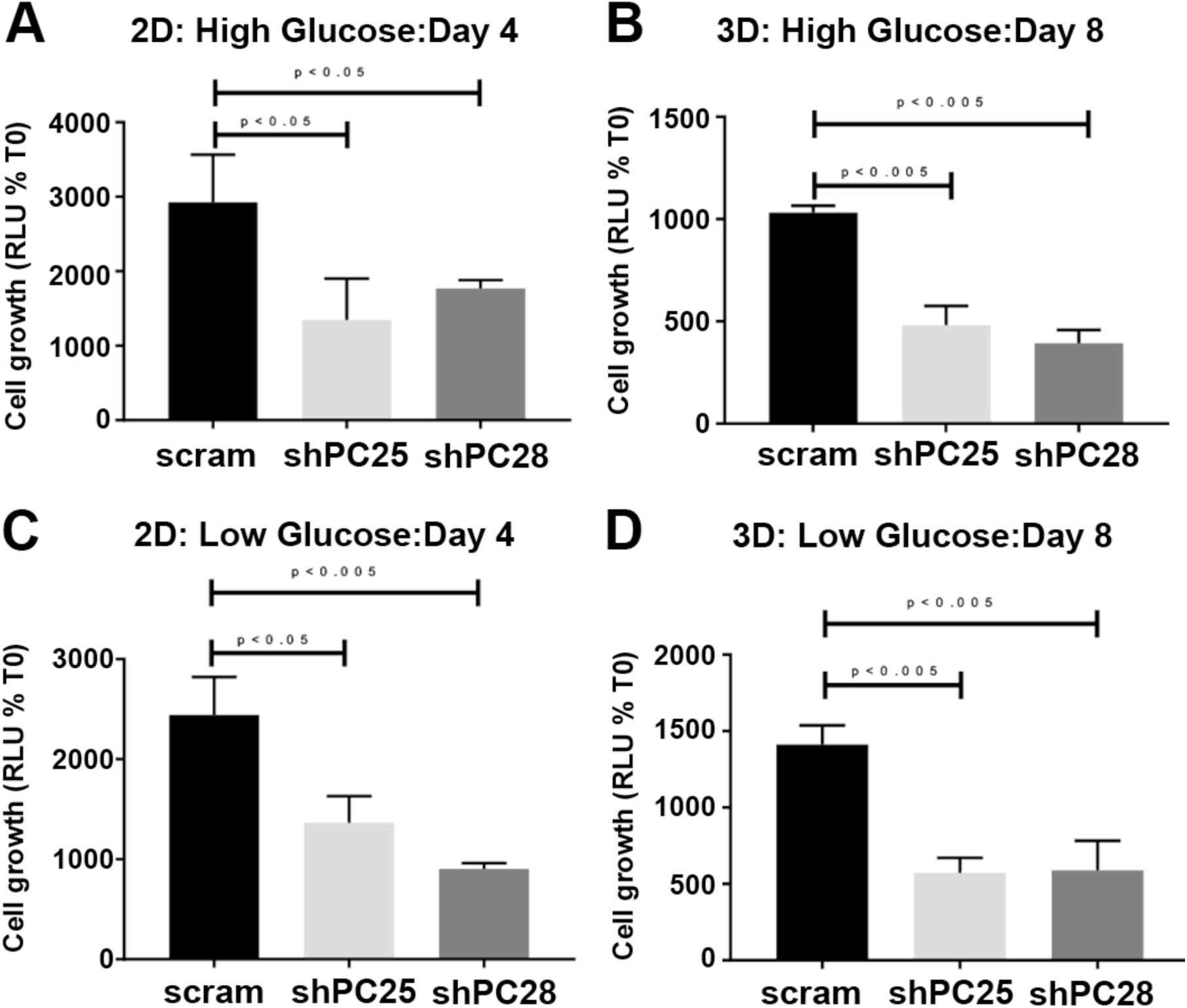
Depletion of the PC inhibits *in vitro* cell growth. Two different shRNA sequences targeting PC (shPC25 and shPC28) were stably expressed in the 4T1 cells and cellular viability was assessed under two-dimensional (A and C) and three-dimensional (B and D) culture conditions containing high (4.5 mg/ml; A and B) or low (1 mg/ml; C and D) amounts of glucose. Data are the mean ± SE of the three separate experiments completed in triplicate resulting in the indicated P values.

**Figure 4.**
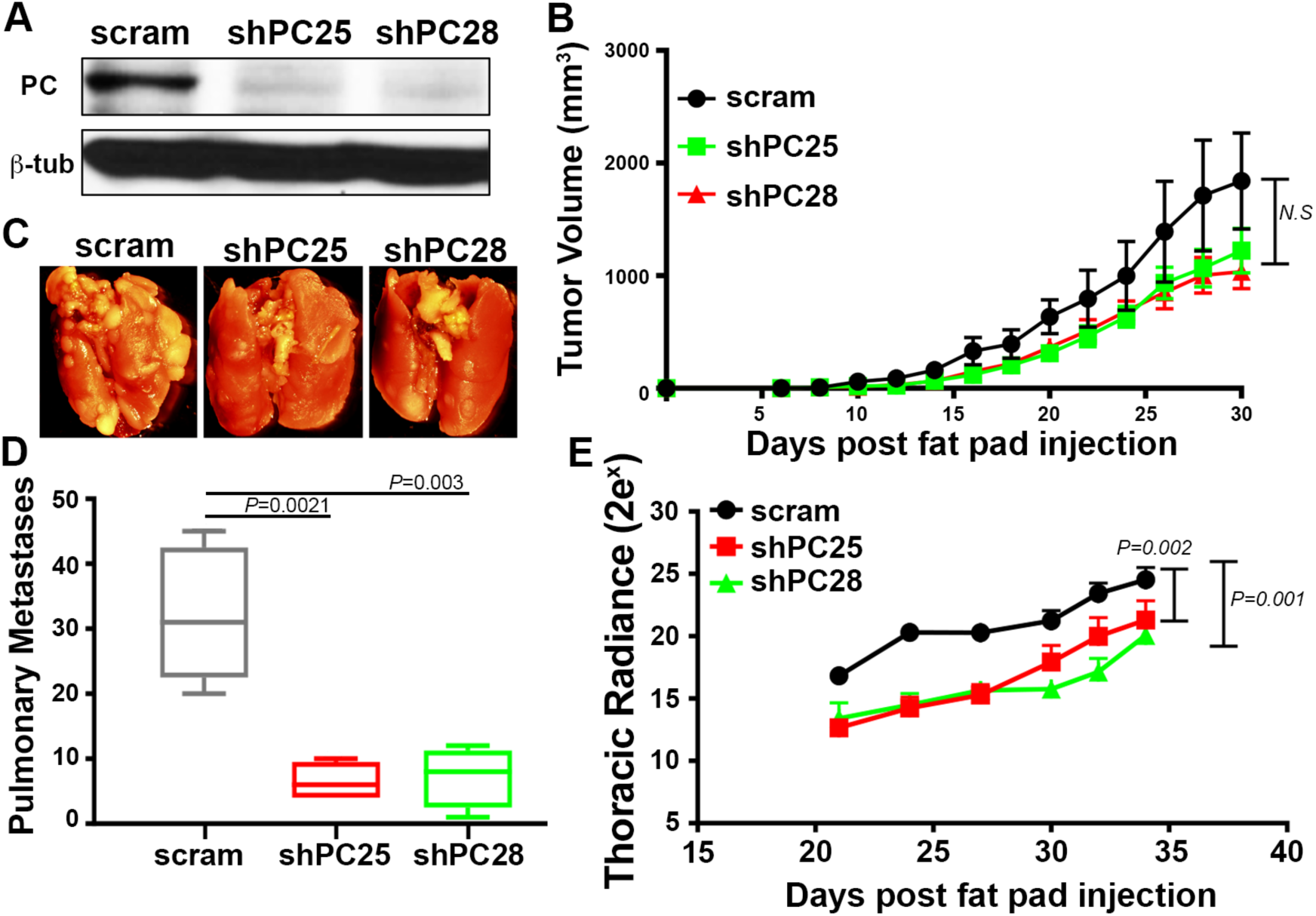
PC is required for 4T1 metastasis but not primary tumor growth. (A) Immunoblot analysis for PC in 4T1 cells expressing control (scram) or PC-targeted (shPC25 and shPC28) shRNAs. Analysis of β-tubulin (β-tub) served as a loading control. (B) 4T1 cells (2.5x10^4^ / mouse) were engrafted onto the mammary fat pad via an intraductal inoculation and primary tumor growth was measured by digital caliper measurements at the indicated time points. Any differences between groups were found to be not significant (N.S.). (C) Necropsy pictures showing lungs of mice bearing the indicated 4T1 primary tumors. (D) The numbers of metastatic pulmonary nodules resulting in mice bearing control (scram) and PC-depleted (shPC25 and shPC28) 4T1 primary tumors. Data are the mean ±SE of five mice per group resulting in the indicated *P* values. (E) Bioluminescent intensity (Radiance) measurements of thoracic metastases of control (scram) or PC-depleted (shPC25 and shPC28) 4T1 tumor bearing mice. Data are the mean radiance measurements ±SE of 5 mice per group resulting the in the indicated *P* values.

### PC expression is required for pulmonary tumor growth

Inhibition of pulmonary metastasis was quite dramatic in the 4T1 model, but overall inhibition of thoracic metastasis, which includes pulmonary metastases, cells within the pleural space and bone metastases throughout the spine and rib cage, was not as robust (Fig. 4d and 4e). These data, together with the lack of change in primary tumor growth (Fig. 4b) and enhanced expression of PC in lung metastases (Fig. 2d) led us to hypothesize that PC may be specifically required for tumor growth within the pulmonary microenvironment. To examine the role of PC in pulmonary tumors we utilized a tail vein injection approach with the D2.A1 cells. These cells grow very well when delivered into the lungs via the tail vein, but do not demonstrate extra-pulmonary metastasis beyond this site [24,25]. Similar to the 4T1 cells PC could be readily depleted from these cells using stable expression of shRNAs (Fig. 5a). Equal pulmonary delivery of control and PC-depleted cells was verified by bioluminescent imaging (Fig. 5b). Furthermore, early bioluminescence readings taken 3 and 8 days after tumor cell injection indicate that initial seeding within in the lungs was not affected by PC depletion (Fig. 5c). However, pulmonary outgrowth was drastically decreased in D2.A1 cells lacking PC (Fig. 5c-5e). These findings strongly suggest that PC is required for initiation of tumor growth within the pulmonary microenvironment.

**Figure 5.**
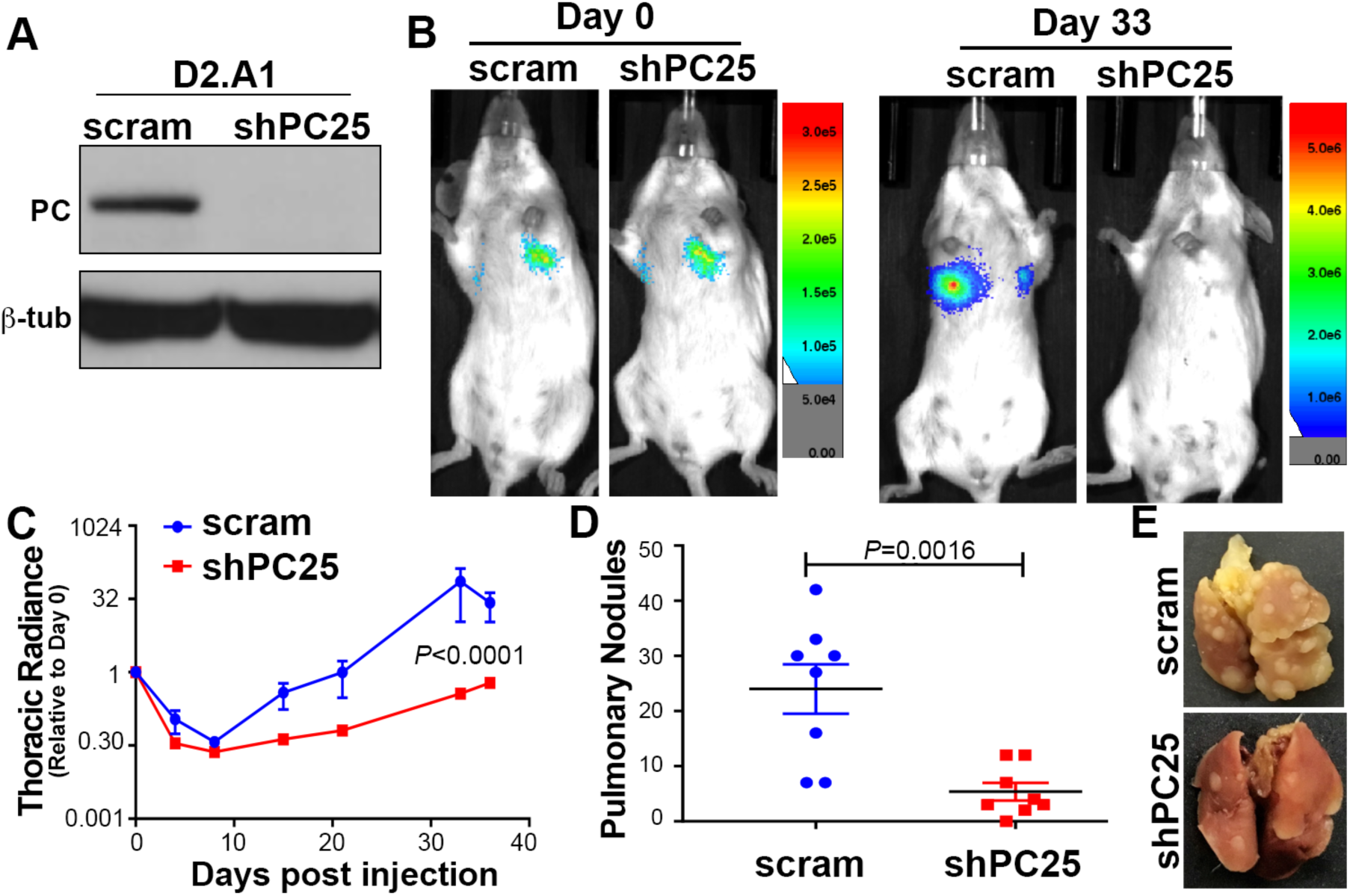
PC expression is required for pulmonary outgrowth of D2.A1 cells. (A) Immunoblot analysis for PC in D2.A1 cells expressing control (scram) or PC-targeted (shPC25) shRNAs. Analysis of β-tubulin (β-tub) served as a loading control. (B) D2.A1 cells (5.0x10^5^ / mouse) were injected into the lateral tail vein of female Balb/c mice. Pulmonary tumor cell delivery (Day 0) and pulmonary tumor growth (Day 33) were visualized by bioluminescent imaging. (C) Bioluminescent intensity (Radiance) values were normalized to the injected values for each group and used to quantify pulmonary tumor seeding and outgrowth at the indicated time points. Growth curves were analyzed two-way ANOVA differences resulting in the indicated P value (n=4 mice per group). (D) The number of pulmonary tumor nodules per lobe resulting in mice injected with control (scram) and PC-depleted (shPC25) D2.A1 cells. Data are the mean ±SE of four mice per group resulting in the indicated *P* values. (E) Necropsy pictures showing representative lungs of mice 35 days after tail vein injections of control (scram) and PC depleted (shPC25) D2.A1 cells.

### PC is not required for growth of extra pulmonary metastases

We have previously established that unlike the D2.A1 cells, the 4T07 cell model will not only grow in the lungs following tail vein inoculation, but will also form extra-pulmonary metastases primarily in the upper thoracic region of the animal [26]. Therefore, we depleted PC in 4T07 cells to evaluate the role of PC in pulmonary versus extra-pulmonary tumor growth following injection into the lateral tail vein (Fig. 6a and 6b). Bioluminescent imaging verified equal pulmonary delivery, but a decrease in ultimate thoracic luminescence generated by PC-depleted 4T07 cells 21 days after tumor cell injection (Fig. 6b and 6c). Upon necropsy a dramatic difference in macroscopic pulmonary tumor nodules once again confirmed the requirement of PC for robust tumor growth within the lungs (Fig. 6d and 6e). In contrast, luminescent imaging of carcasses in which tumor-bearing lungs had been removed demonstrated a significant increase in extra-pulmonary metastases generated by PC-depleted cells as compared to control cells (Fig. 6f and 6g). These data strongly suggest that while PC is required for growth in the lungs its expression is not required, or maybe inhibitory to the growth of extra-pulmonary metastases.

**Figure 6.**
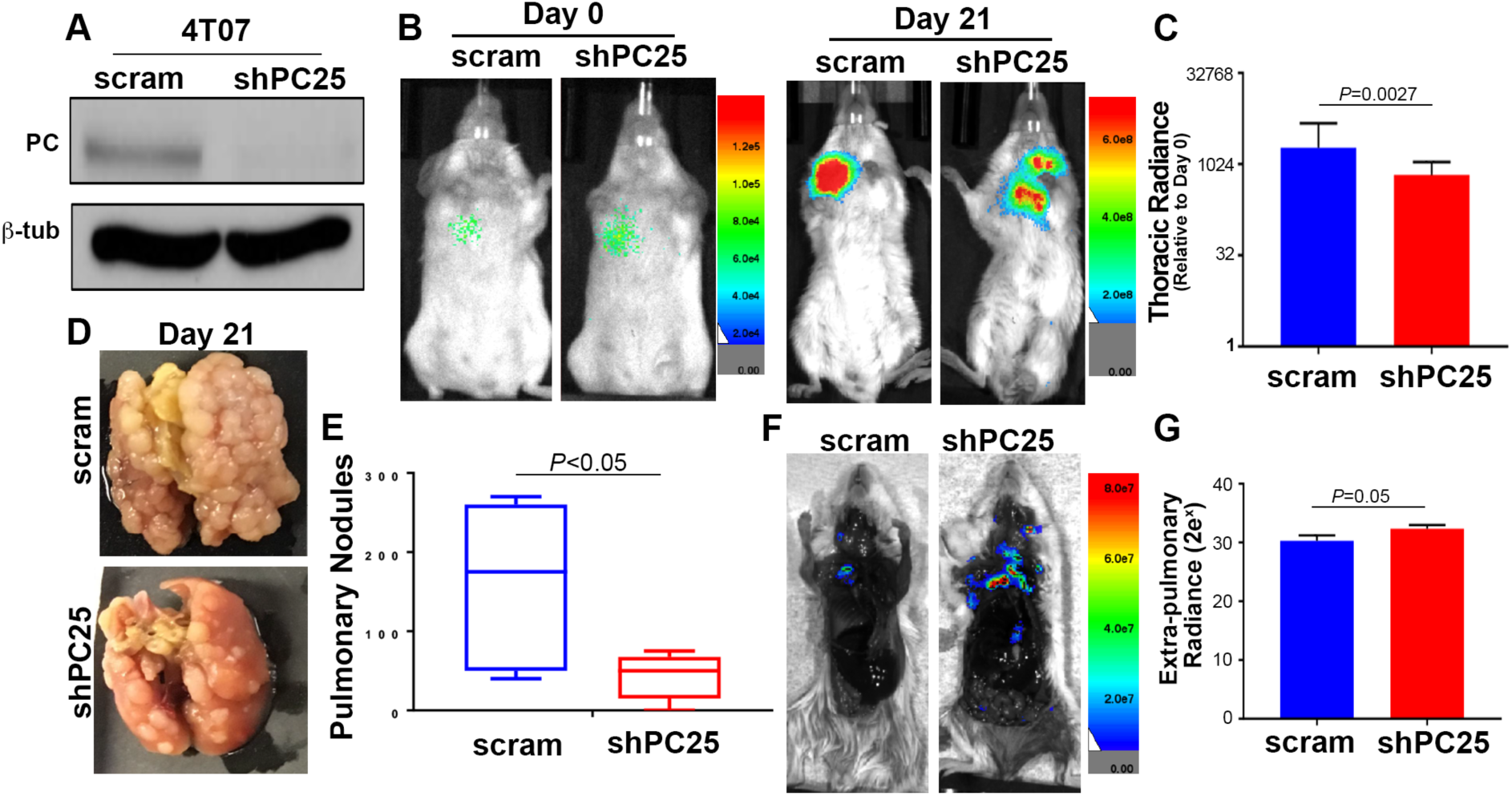
PC is not required for the growth of extra pulmonary metastases. (A) Immunoblot analysis for PC in 4T07 cells expressing control (scram) or PC-targeted (shPC25) shRNAs. Analysis of β-tubulin (β-tub) served as a loading control. (B) 4T07 cells (5.0x10^5^ / mouse) were injected into the lateral tail vein of female Balb/c mice. Pulmonary tumor cell delivery (Day 0) and tumor formation (Day 21) were visualized by bioluminescent imaging. (C) Bioluminescent intensity (Radiance) values were normalized to the injected values for each group and used to quantify thoracic tumor growth in intact mice 21 days after tumor cell injections. Data are the mean normalized values ±SE of five mice per group resulting in the indicated *P* value. (D) Necropsy pictures showing representative lungs of mice 21 days after tail vein injections of control (scram) and PC depleted (shPC25) 4T07 cells. (E) The numbers of pulmonary tumor nodules per lobe was quantified in mice injected with control (scram) and PC-depleted (shPC25) 4T07 cells. Data are the mean ±SE of five mice per group resulting in the indicated *P* value. (F) The lungs of mice injected with control (scram) and PC-depleted (shPC25) 4T07 cells were removed upon necropsy and the carcasses were immediately imaged. Shown are representative images from each group. (G) Whole animal bioluminescent intensity measurements were taken for dissected carcasses described in panel F. Data are the mean radiance values ±SE of five mice per group resulting in the indicated *P* value.

### PC is required for the metabolic plasticity of metastatic breast cancer

To investigate the importance of PC in the metabolic plasticity of metastatic breast cancer cells, rates of glycolysis and oxygen consumption were measured using the XFe24 Seahorse bioanalyzer. Both glycolysis, as measured by the extracellular acidification rate (ECAR), and oxygen consumption rate (OCR) were significantly higher in 4T1 versus 4T07 cells (Fig. 7; [7]). Along these lines, PC depletion resulted in a significant decrease in both glycolysis and OCR in 4T1 cells, but not 4T07 cells (Fig 7a and 7c). Glycolytic capacity was further measured by injecting oligomycin (Fig. 7b), an ATP synthase inhibitor which shifts cell dependence on ATP generation towards glycolysis. Consistent with previous reports, 4T1 cells demonstrated significantly higher glycolytic capacity than the 4T07 cells, and again this was dependent upon expression of PC (Fig 7a; [27]).

**Figure 7.**
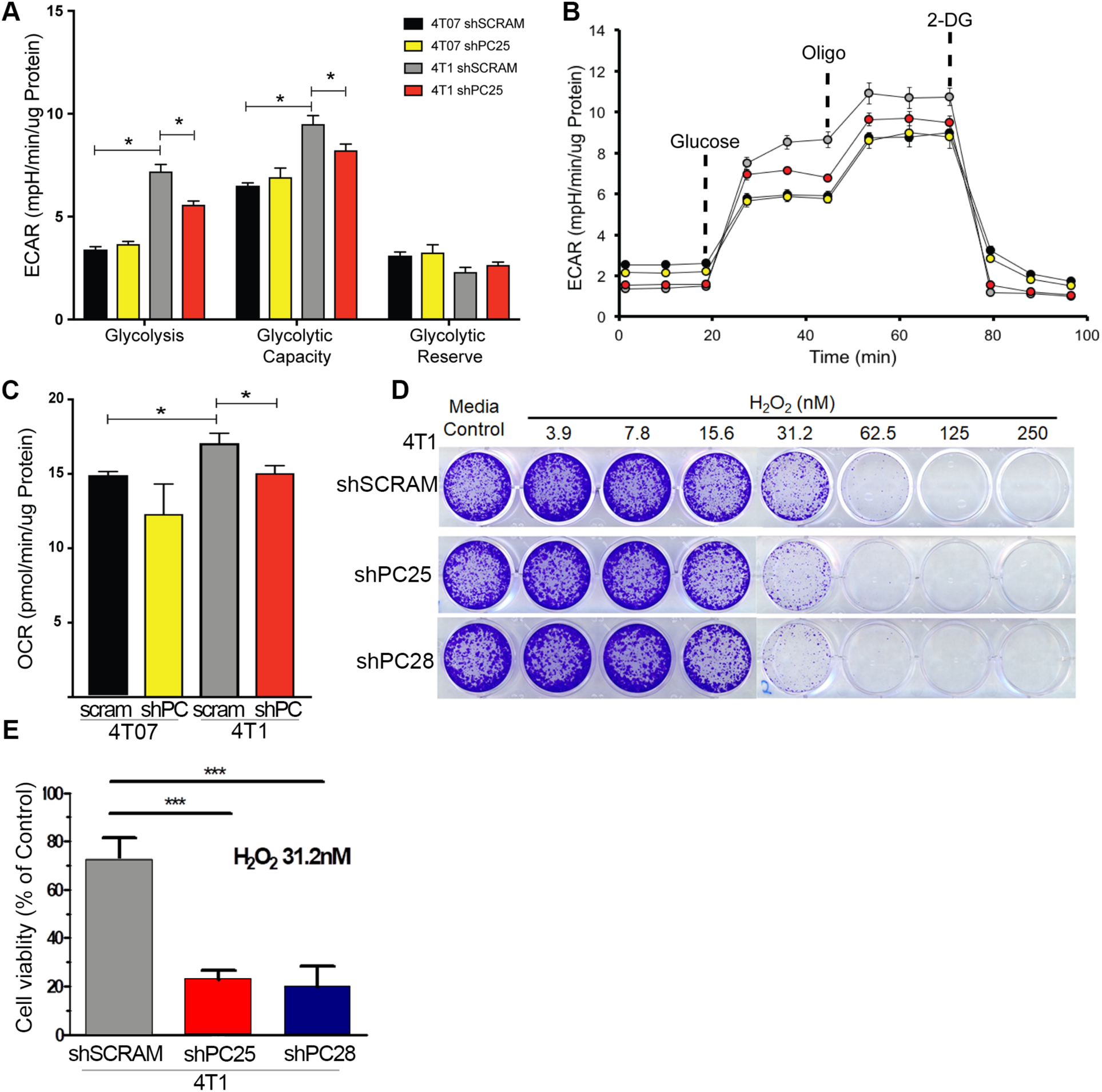
PC supports metabolic flexibility and oxidative stress defense. (A) Glycolytic stress test analysis of 4T07 and 4T1 control (shscram) and PC depleted (shPC25) cells. (B) ECAR measurements obtained upon injections of glucose (10mM), oligomycin (Oligo) and 2-deoxglucose (2-DG) used to calculate glycolysis and glycolytic capacity in each cell line. (C) OCR measurements for 4T07 and 4T1 control (shscram) and PC depleted (shPC25) cells in the presence of 10 mM glucose, 2mM glutamine and 1mM pyruvate. Data are the mean ±SE of measurements per μg protein. (D) Control (shscram) and PC-depleted (shPC25 and shPC28) 4T1 cells were treated with the indicated concentrations of H_2_O_2_ and viable cells were visualized with crystal violet. (E) Stained cells from panel D were solubilized and quantified via absorbance at 600nm. Data are the mean ±SD of three independent experiments. Asterisks indicates statistical significance (*<0.05, ***<0.01).

The hypoxic environment of the primary tumor induces the stabilization of HIF-1a and shut down of the TCA cycle, but newly developing pulmonary metastases lose HIF-1α expression (Figure S5; [28]). Thus, metabolic adaptation upon loss of HIF-1α is a major hurdle that must be overcome during initiation of pulmonary metastatic outgrowth. HIF-1α levels are constitutively low in *in vitro* culture, consistent with the requirement of PC for *in vitro* cell growth (Fig 3). Therefore, to further recapitulate the enhanced oxidative stress within the pulmonary microenvironment we treated cells with increasing concentrations of hydrogen peroxide (H_2_O_2_). Consistent with our recent studies we observed an enhanced sensitivity to H_2_O_2_ upon depletion of PC (Fig 7d and 7e; [29]).

## Discussion

Herein, we utilized several syngeneic models of breast cancer to demonstrate that initiation of pulmonary metastatic outgrowth is strongly dependent on PC. We also demonstrate that PC is not required for primary mammary tumor growth or extra-pulmonary metastasis. Our findings are nicely supported by previous studies in lung cancer that similarly demonstrate the requirement of PC for *in vivo* tumor growth [10,14]. The specific aspects of the pulmonary microenvironment that demand PC expression remain to be determined definitively. However, enhanced oxygen content and increased oxidative stress are very unique aspects of the pulmonary microenvironment with respect to pyruvate utilization. Under hypoxic conditions of primary tumors, extrapulmonary metastases, and macroscopic pulmonary tumors, pyruvate utilization mainly occurs through anaerobic glycolysis that is driven by HIF-1α-induced expression of hexokinase, LDH, and pyruvate kinase M2 [28]. HIF-1α also inhibits pyruvate dehydrogenase via upregulation of PDK1, further blocking aerobic glycolysis and increasing lactate production [30]. Depletion of HIF-1α or HIF-1α target genes such as LDH or carbonic anhydrase IX (CAIX) in 4T1 cells inhibits primary tumor growth, therefore decreasing subsequent macrometastasis [12,31][13]. In contrast, we propose a model that in the oxygen and pyruvate rich microenvironment of the lungs HIF-1α is directed to proteosomal degradation and metastatic cells must transiently shift to PC-mediated aerobic utilization of pyruvate to initiate metastatic outgrowth. Only once macroscopic metastases are formed and hypoxia is reestablished are tumors able to reengage HIF-1α and return to anaerobic glycolysis.

Our analysis of the METABRIC dataset demonstrate decreased survival of stage 1 breast cancer patients with genomic amplification of PC. These findings are completely consistent with our mouse studies in that high-level PC expression does not contribute to primary tumor growth, but these cells have an enhanced antioxidant defense system and are more metabolically fit for aerobic initiation of pulmonary metastatic outgrowth following systemic dissemination. Indeed, recent studies clearly indicate that systemic dissemination of primary tumor cells is an early step in metastasis whereas overcoming dormancy within secondary organs is the rate limiting process leading to patient lethality [32]. Therefore, our findings suggest assessment of PC genomic amplification or mRNA expression could serve as an effective prognostic biomarker to predict for metastatic relapse in patients diagnosed with early stage primary tumors. Additionally, PC could also serve as therapeutic biomarker in conjunction with application of direct pharmacological inhibitors of PC or more general inhibitors of oxidative phosphorylation such as IACS-010759, which is currently in clinical trials (NCT03291938).

## Conclusions

Overall, the data herein present a comprehensive analysis of patient data and syngeneic mouse models that uniformly support a specific role for PC in facilitating initiation of pulmonary metastatic tumor growth. Moreover, we present findings that demonstrate a role for PC in facilitating metabolic plasticity of disseminated breast cancer cells. Our findings clearly point to the potential of genomic amplification of PC as prognostic biomarker and as a therapeutic target for the treatment of pulmonary metastatic breast cancer.

**List of abbreviations**
PC: – Pyruvate carboxylase
EMT: – epithelial-mesenchymal transition
HIF1-α: - Hypoxia Inducible factor
METABRIC: - Molecular Taxonomy of Breast Cancer International Consortium
MTCI: - Molecular Therapeutics of Cancer, Ireland
HAN: - Hyperplastic Alveolar Nodule
LDH: - Lactate Dehydrogenase
OXPHOS: – Oxidative phosphorylation
PGC-1α: - Peroxisome proliferator-activated receptor gamma coactivator 1-alpha
PDH: – Pyruvate Dehydrogenase
PDK1: – Pyruvate Dehydrogenase kinase
OCR: – Oxygen consumption rate
ECAR: – Extra Cellular Acidification Rate

## Declarations

**Ethics approval and consent to participate:** All animal studies were performed in accordance with the animal protocol procedures approved by the Purdue Animal Care and Use Committee of Purdue University.

**Consent for publication:** Not applicable

**Availability of data and materials:** All data generated or analyzed during this study are included in this published article and its supplementary information files.

**Competing interests:** The authors declare no competing interests.

**Funding:** This research was supported in part by the American Cancer Society (RSG-CSM130259) to M.K.W. and the National Institutes of Health (R01CA207751) to M.K.W., the National Institutes of Health (R25CA128770) to DT and the Indiana Elks Charities to D.T. Support was also provide by the Indiana Clinical and Translational Science Institute to M.K.W. and D.T. (UL1TR001108) and the Purdue Center for Cancer Research via an NIH NCI grant (P30CA023168).

## Author Contributions

A.S., H.C. and T.W. contributed to the design of the experiments, completed experiments and contributed to the writing of the manuscript. M.K.W. and D.T. contributed to the design of the experiments, supervised experiments and contributed to the writing of the manuscript.

## Acknowledgments

Members of the Wendt Laboratory are thanked for critical reading of the manuscript. We kindly acknowledge the expertise of the personnel within the Purdue Center for Cancer Research Biological Evaluation Core. We also acknowledge the use of the facilities within the Bindley Bioscience Center, a core facility of the NIH-funded Indiana Clinical and Translational Sciences Institute.

## Supplementary Figures

**Figure S1.**
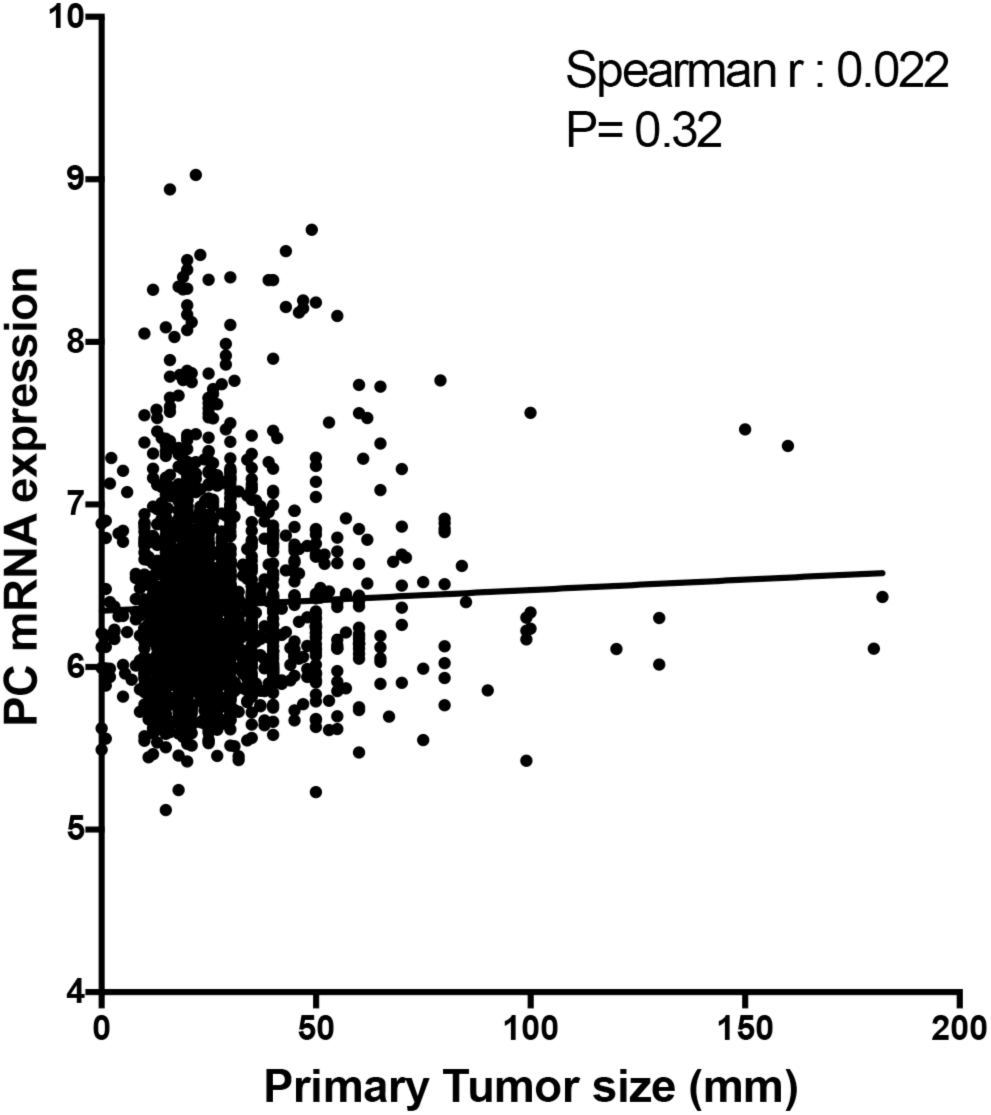
PC expression is not correlated with primary tumor size. Patient samples within the METABRIC dataset were analyzed for PC expression in relation to primary tumor size. Data are analyzed by a non-parametric spearman correlation resulting in the indicated r and P values. The linear regression for these data is also shown.

**Figure S2.**
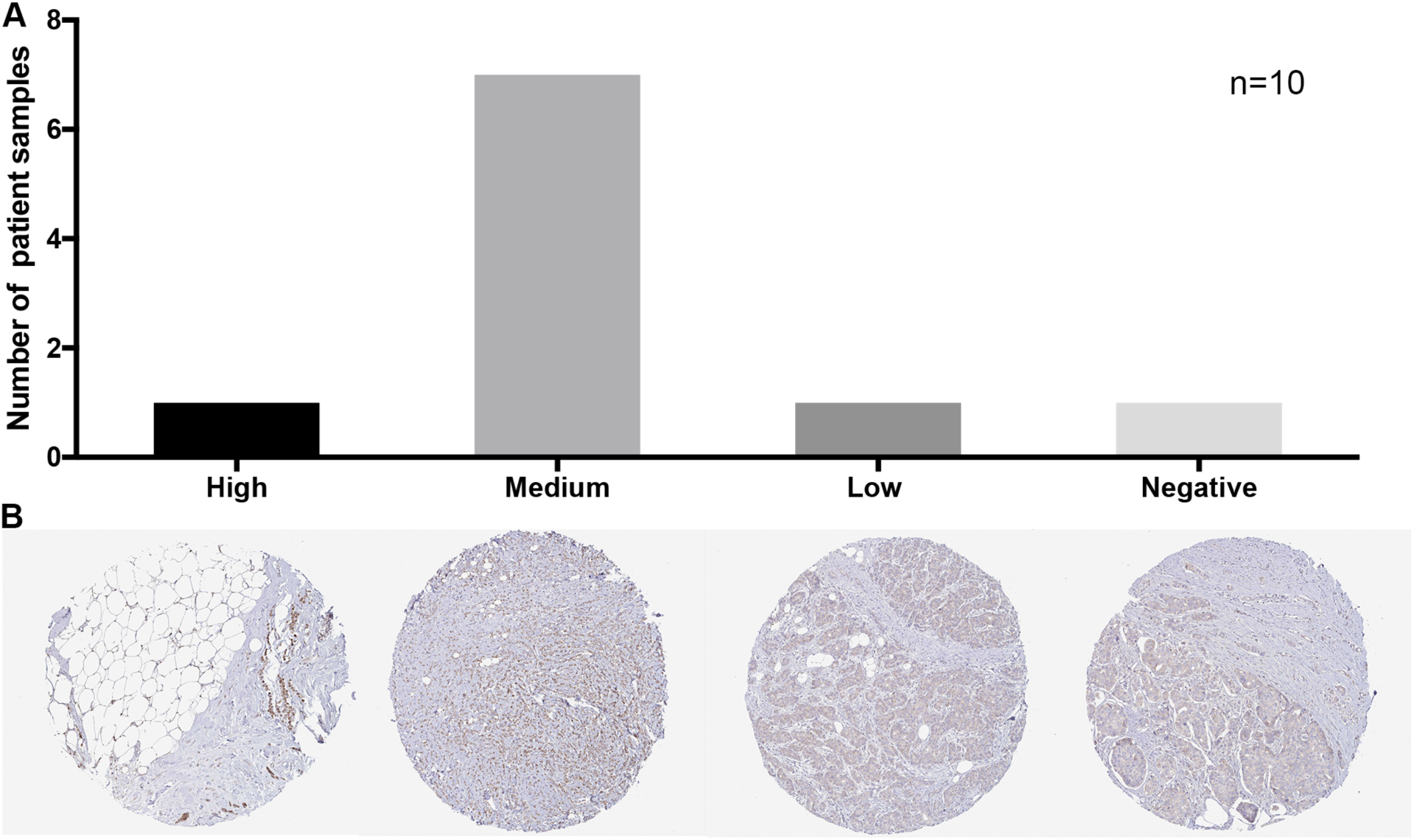
PC expression in primary breast tumors. PC expression within tumor cells was rated as high, medium, low or negative (n=10). Representative sections from each group are shown. Data was obtained from the protein atlas dataset (www.proteinatlas.org) [33].

**Figure S3.**
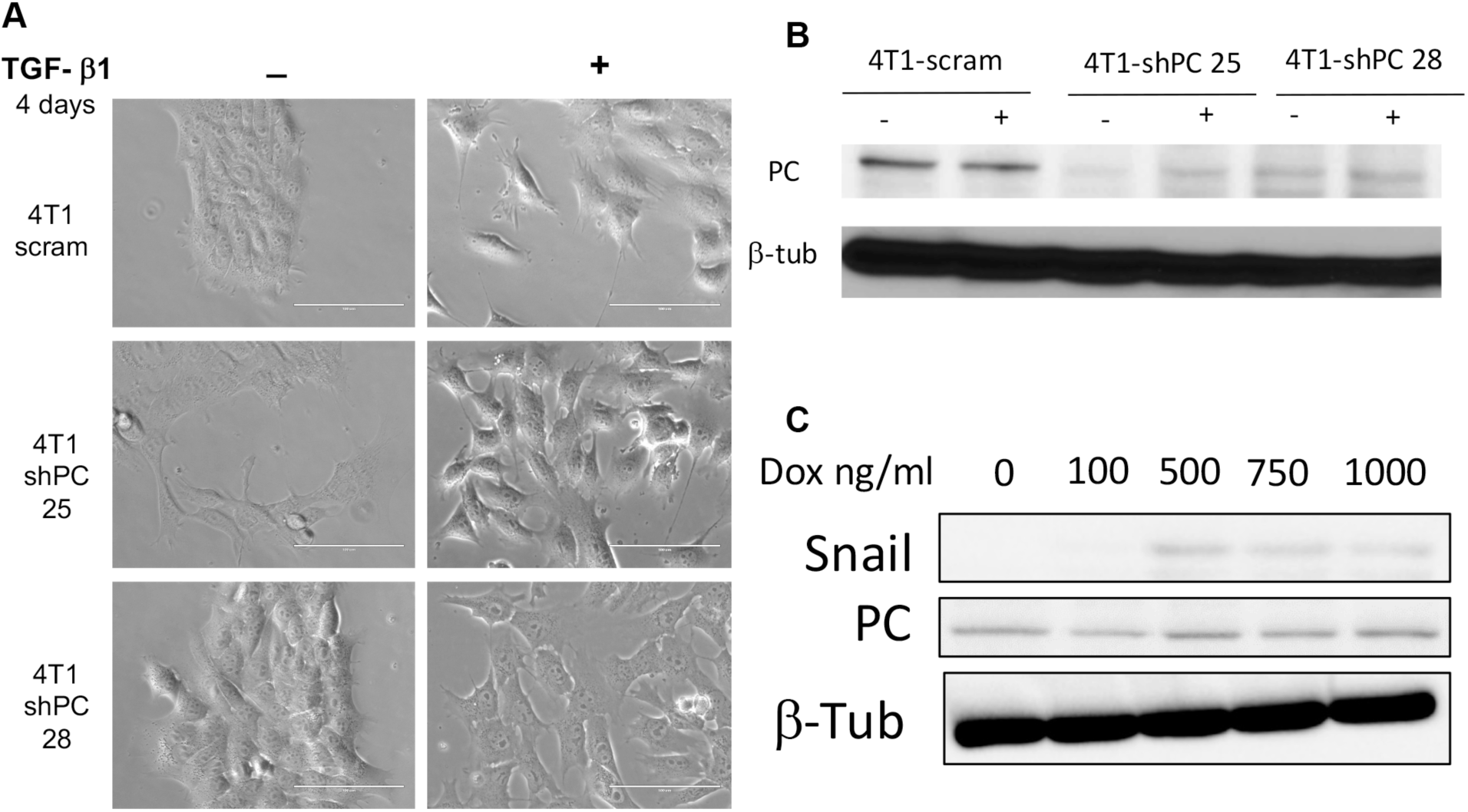
Transient induction of EMT does not induce expression of PC. (A) Photomicrographs showing EMT-like changes in cellular morphology of the control (scram) and PC-depleted (shPC25 and shPC28) 4T1 cells upon treatment with exogenous TGF-β1 (5ng/ml) for 4 days. (B) Immunoblot analyses for PC in cells shown in Panel A. β-tubulin served as a loading control. (C) The RAS transformed MCF-10A-T1k cells were constructed to express a doxycycline-inducible vector encoding the EMT transcription factor Snail. Expression of PC was analyzed upon addition of Doxycycline (Dox) at the indicated concentrations. Expression of Snail and β-tubulin (β-Tub) served as loading controls.

**Figure S4.**
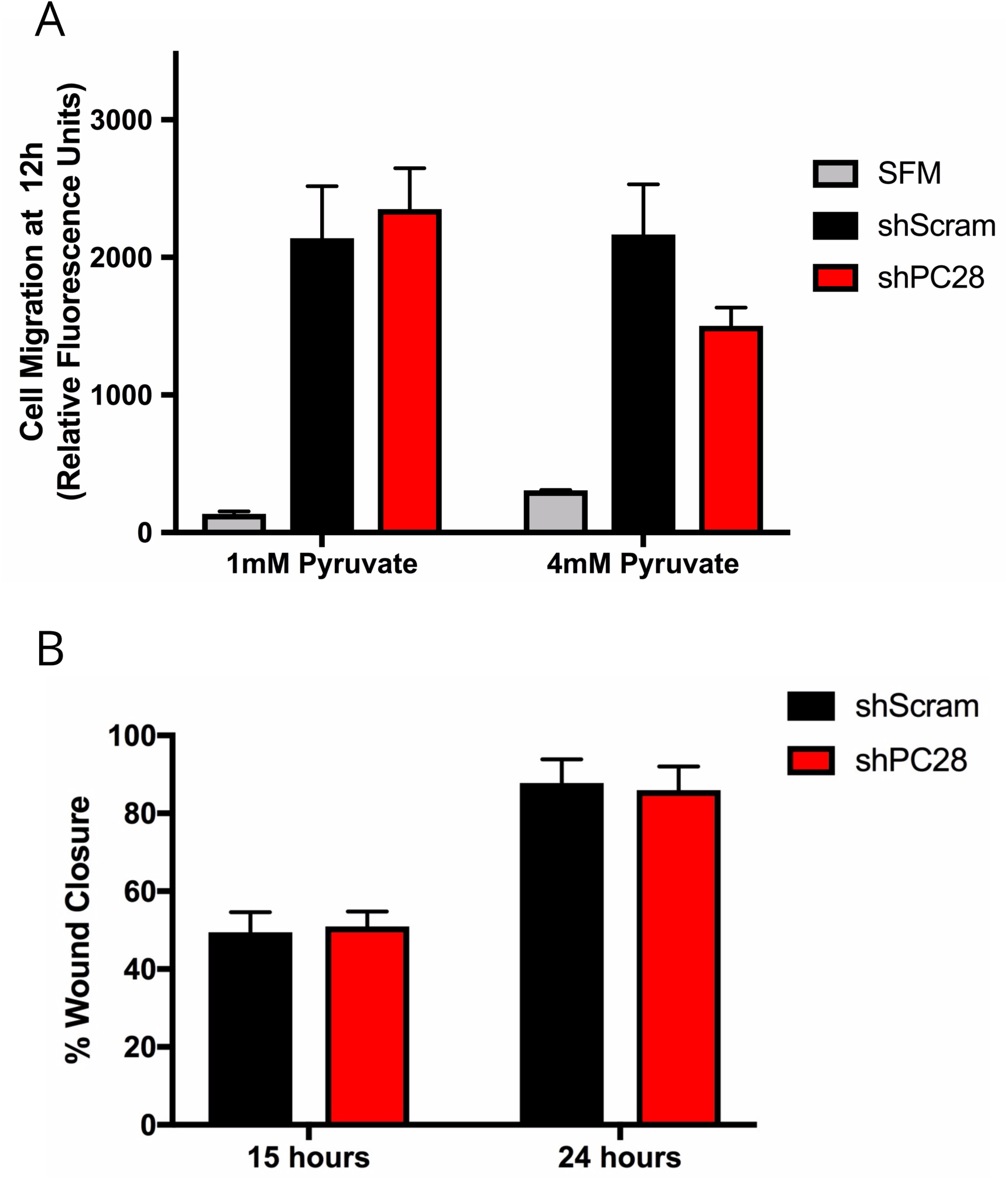
PC is not required for cell migration. (A) Control (shScram) and PC depleted (shPC28) cells were plated into 8μm transwell inserts in serum free media containing 1 or 4 mM pyruvate. Cell migration was quantified 12 hours after plating using Calcein AM. SFM represents wells where the bottom chamber was filled with serum free media as negative control. Values are presented as mean relative fluorescence units, ±S.E.M. (B) Control (shScram) and PC depleted (shPC28) 4T1 monolayers were wounded and closure was measured 15 and 24 hours later. Values are presented as mean percent would closure, ±S.E.M.

**Figure S5.**
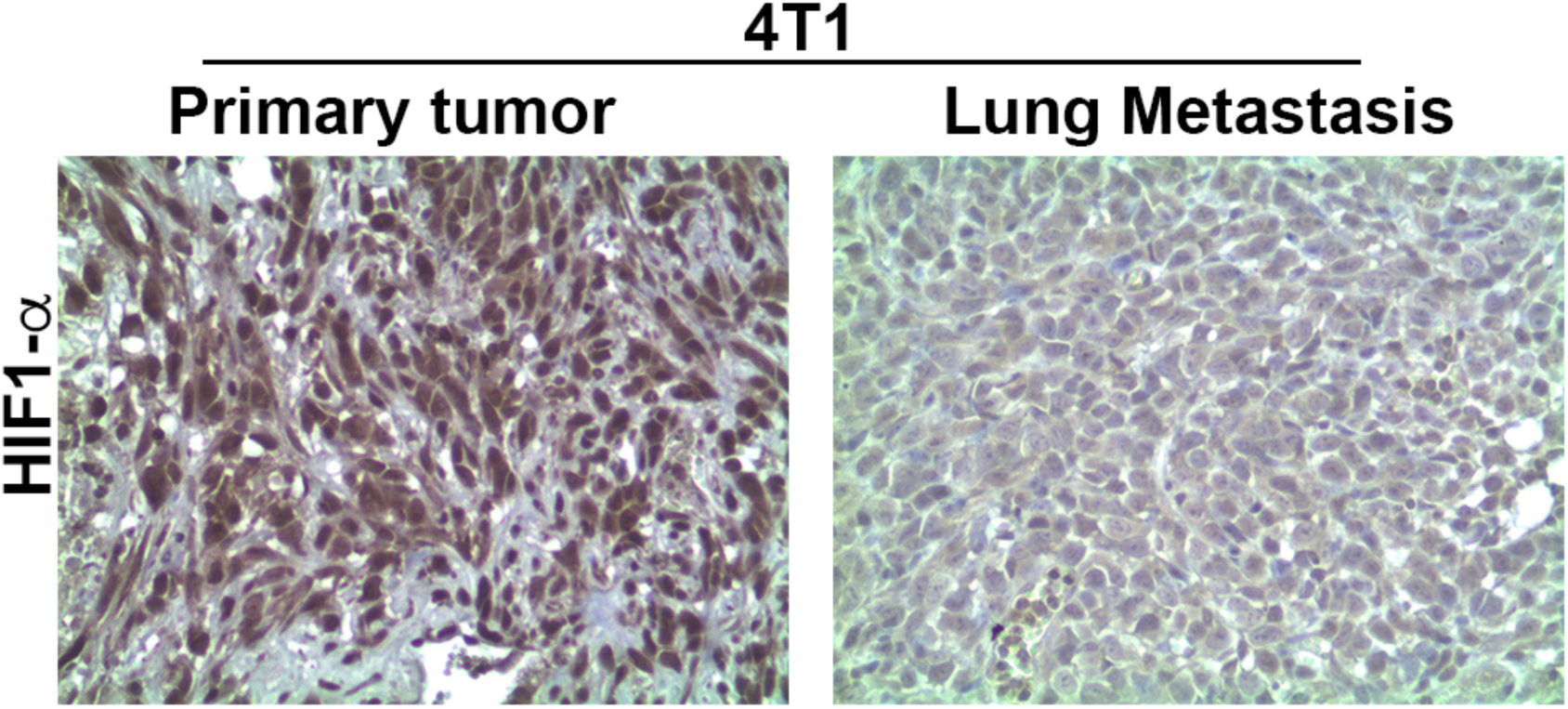
HIF-1α expression that characterizes primary tumors is lost in pulmonary metastases. 4T1 cells were engrafted onto the mammary fat pad via an intraductal injection and grown as primary tumors. These tumors gave rise to spontaneous pulmonary metastases. Upon necropsy both primary and metastatic tumors were analyzed by immunohistochemistry for the expression of HIF-1α. Nuclear expression of HIF-1α is very high in viable primary tumor tissue. In contrast, nuclear HIF-1α is drastically reduced in pulmonary metastases. Data are representative of 3 separate mice bearing primary tumors and metastases.

## References

1. Ali R, Wendt MK. The paradoxical functions of EGFR during breast cancer progression. Signal Transduct Target Ther. 2017;2:16042.

2. Chambers AF, Groom AC, MacDonald IC. Dissemination and growth of cancer cells in metastatic sites. Nat Rev Cancer. 2002;2:563–72.

3. Minn AJ, Kang Y, Serganova I, Gupta GP, Giri DD, Doubrovin M, et al. Distinct organ-specific metastatic potential of individual breast cancer cells and primary tumors. J Clin Invest. 2005;115:44–55.

4. Hoshino A, Costa-Silva B, Shen T-L, Rodrigues G, Hashimoto A, Tesic Mark M, et al. Tumour exosome integrins determine organotropic metastasis. Nature. 2015;527:329–35.

5. Hanahan D, Weinberg RA. Hallmarks of Cancer: The Next Generation. Cell. 2011;144:646–74.

6. Heiden MGV, DeBerardinis RJ. Understanding the Intersections between Metabolism and Cancer Biology. Cell. 2017;168:657–69.

7. Simões RV, Serganova IS, Kruchevsky N, Leftin A, Shestov AA, Thaler HT, et al. Metabolic Plasticity of Metastatic Breast Cancer Cells: Adaptation to Changes in the Microenvironment. Neoplasia. 2015;17:671–84.

8. Andrzejewski S, Klimcakova E, Johnson RM, Tabariès S, Annis MG, McGuirk S, et al. PGC-1α Promotes Breast Cancer Metastasis and Confers Bioenergetic Flexibility against Metabolic Drugs. Cell Metab. 2017;26:778–787.e5.

9. Phannasil P, Thuwajit C, Warnnissorn M, Wallace JC, MacDonald MJ, Jitrapakdee S. Pyruvate Carboxylase Is Up-Regulated in Breast Cancer and Essential to Support Growth and Invasion of MDA-MB-231 Cells. PloS One. 2015;10:e0129848.

10. Sellers K, Fox MP, Bousamra M, Slone SP, Higashi RM, Miller DM, et al. Pyruvate carboxylase is critical for non-small-cell lung cancer proliferation. J Clin Invest. 2015;125:687–98.

11. Christen S, Lorendeau D, Schmieder R, Broekaert D, Metzger K, Veys K, et al. Breast Cancer-Derived Lung Metastases Show Increased Pyruvate Carboxylase-Dependent Anaplerosis. Cell Rep. 2016;17:837–48.

12. Rizwan A, Serganova I, Khanin R, Karabeber H, Ni X, Thakur S, et al. Relationships between LDH-A, Lactate, and Metastases in 4T1 Breast Tumors. Clin Cancer Res. 2013;19:5158–69.

13. Dupuy F, Tabariès S, Andrzejewski S, Dong Z, Blagih J, Annis MG, et al. PDK1-Dependent Metabolic Reprogramming Dictates Metastatic Potential in Breast Cancer. Cell Metab. 2015;22:577–89.

14. Davidson SM, Papagiannakopoulos T, Olenchock BA, Heyman JE, Keibler MA, Luengo A, et al. Environment Impacts the Metabolic Dependencies of Ras-Driven Non-Small Cell Lung Cancer. Cell Metab. 2016;23:517–28.

15. Gebäck T, Schulz MMP, Koumoutsakos P, Detmar M. TScratch: a novel and simple software tool for automated analysis of monolayer wound healing assays. BioTechniques. 2009;46:265–74.

16. Madden SF, Clarke C, Gaule P, Aherne ST, O’Donovan N, Clynes M, et al. BreastMark: an integrated approach to mining publicly available transcriptomic datasets relating to breast cancer outcome. Breast Cancer Res BCR. 2013;15:R52.

17. Pereira B, Chin S-F, Rueda OM, Vollan H-KM, Provenzano E, Bardwell HA, et al. The somatic mutation profiles of 2,433 breast cancers refines their genomic and transcriptomic landscapes. Nat Commun. 2016;7:11479.

18. Strickland LB, Dawson PJ, Santner SJ, Miller FR. Progression of premalignant MCF10AT generates heterogeneous malignant variants with characteristic histologic types and immunohistochemical markers. Breast Cancer Res Treat. 2000;64:235–40.

19. Morris VL, Tuck AB, Wilson SM, Percy D, Chambers AF. Tumor progression and metastasis in murine D2 hyperplastic alveolar nodule mammary tumor cell lines. Clin Exp Metastasis. 1993;11:103–12.

20. Aslakson CJ, Miller FR. Selective events in the metastatic process defined by analysis of the sequential dissemination of subpopulations of a mouse mammary tumor. Cancer Res. 1992;52:1399–405.

21. Lee SY, Jeon HM, Ju MK, Kim CH, Yoon G, Han SI, et al. Wnt/Snail Signaling Regulates Cytochrome c Oxidase and Glucose Metabolism. Cancer Res. 2012;72:3607–17.

22. Jitrapakdee S, Maurice MS, Rayment I, Cleland WW, Wallace JC, Attwood PV. Structure, Mechanism and Regulation of Pyruvate Carboxylase. Biochem J. 2008;413:369–87.

23. Kumashiro N, Beddow SA, Vatner DF, Majumdar SK, Cantley JL, Guebre-Egziabher F, et al. Targeting pyruvate carboxylase reduces gluconeogenesis and adiposity and improves insulin resistance. Diabetes. 2013;62:2183–94.

24. Wendt MK, Taylor MA, Schiemann BJ, Schiemann WP. Down-regulation of epithelial cadherin is required to initiate metastatic outgrowth of breast cancer. Mol Biol Cell. 2011;22:2423–35.

25. Shibue T, Weinberg RA. Integrin beta1-focal adhesion kinase signaling directs the proliferation of metastatic cancer cells disseminated in the lungs. Proc Natl Acad Sci U S A. 2009;106:10290–5.

26. Wendt MK, Schiemann BJ, Parvani JG, Lee Y-H, Kang Y, Schiemann WP. TGF-β stimulates Pyk2 expression as part of an epithelial-mesenchymal transition program required for metastatic outgrowth of breast cancer. Oncogene. 2013;32:2005–15.

27. Frees AE, Rajaram N, McCachren SS, Fontanella AN, Dewhirst MW, Ramanujam N. Delivery-corrected imaging of fluorescently-labeled glucose reveals distinct metabolic phenotypes in murine breast cancer. PloS One. 2014;9:e115529.

28. Kim J, Tchernyshyov I, Semenza GL, Dang CV. HIF-1-mediated expression of pyruvate dehydrogenase kinase: a metabolic switch required for cellular adaptation to hypoxia. Cell Metab. 2006;3:177–85.

29. Wilmanski T, Zhou X, Zheng W, Shinde A, Donkin SS, Wendt M, et al. Inhibition of pyruvate carboxylase by 1α,25-dihydroxyvitamin D promotes oxidative stress in early breast cancer progression. Cancer Lett. 2017;411:171–81.

30. Kim J, Gao P, Liu Y-C, Semenza GL, Dang CV. Hypoxia-inducible factor 1 and dysregulated c-Myc cooperatively induce vascular endothelial growth factor and metabolic switches hexokinase 2 and pyruvate dehydrogenase kinase 1. Mol Cell Biol. 2007;27:7381–93.

31. Lou Y, McDonald PC, Oloumi A, Chia S, Ostlund C, Ahmadi A, et al. Targeting tumor hypoxia: suppression of breast tumor growth and metastasis by novel carbonic anhydrase IX inhibitors. Cancer Res. 2011;71:3364–76.

32. Sosa MS, Bragado P, Aguirre-Ghiso JA. Mechanisms of disseminated cancer cell dormancy: an awakening field. Nat Rev Cancer. 2014;14:611–22.

33. Pontén F, Jirström K, Uhlen M. The Human Protein Atlas--a tool for pathology. J Pathol. 2008;216:387–93.

